# “Homogenous subgroups of atypical meningiomas defined using oncogenic signatures: basis for a new grading system?”

**DOI:** 10.1101/493031

**Authors:** Zsolt Zador, Alexander Landry, Ashirbani Saha, Michael D. Cusimano

## Abstract

Meningiomas are the most common brain tumor with a prevalence of 3% in the population. Histological grading of meningiomas (1 through 3) has a major role in determining treatment choice and predicting outcome. While largely indolent grade 1 and the highly aggressive grade 3 meningiomas as considered mostly homogenous in clinical behavior, atypical or grade 2 meningiomas have highly diverse biological properties. Our aim was to identify homogenous subgroups of atypical meningiomas with the working hypothesis that these subgroups would share features with grade 1 and grade 3 counterparts. We carried out systems level analysis by gene module discovery using co-expression networks on the transcriptomics of 212 meningiomas. The newly identified subgroups were characterized in terms of recurrence rate and overlapping biological processes in gene ontology. We were able to reclassify 33 of 46 atypical meningiomas (72%) into a benign “grade 1-like” (14/46) and malignant “grade 3-like” (19/46) subgroup based on oncogenic signatures. Recurrence rates of “Grade 1-like” and “grade 3-like” tumors was 0% and 72% respectively. These two new subgroups showed similar recurrence rates and concordant biological processes with the respected grades. Our findings help resolve the heterogeneity/uncertainty around atypical meningioma biology and identify subgroups more homogenous than in prior studies. These results may help reshape prediction, follow-up planning, treatment decisions and recruitment protocols for future and ongoing clinical trials. The findings demonstrate the conceptual advantage of systems biology approaches and underpin the utility of molecular signatures as complements to the current histological grading system.

## Introduction

Meningiomas are the most common adult brain tumor, carrying an overall prevalence of 3% in the population. Histopathologic analysis is the mainstay of diagnosis and together with the extent of surgical resection it is a determinant of outcome and treatment planning^1,2^. According to World Health Organization (WHO) grading, majority of meningiomas (almost 70%) fall into grade 1, of which about two thirds are cured with surgical excision alone^2^ and 15-20 % recur within five years of diagnosis^3^. Grade 3 meningiomas, by contrast, are rare and aggressive with a five year recurrence rate of approximately 90%^2^. Grade 2 (atypical) meningiomas constitute 20-30% of cases and represent a biological intermediate. Predicting the clinical course for these tumors is particularly challenging given their heterogeneous biology which yields a five-year recurrence rate of approximately 50%^4^. Previous studies have suggested that some grade 2 meningiomas share features with grade 1s while others are more similar to grade 3s based on clinical behavior as well as genetic features such as somatic mutations, copy number variants^5–7^, methylation status^8^, and genome wide expression profiles^9,10^. However, most research on gene expression in meningioma focuses on single gene analytics. This is not optimized for the low and additive molecular signals which frequently underlie complex and heterogeneous diseases. Systems biology approaches such as coexpression networks^11,12^, on the other hand, are able to provide a higher resolution of these complex genetic processes^11,13–15^. We therefore hypothesized that such methods may reveal subgroups for grade 2 meningiomas based on resemblance to grade 1 or grade 3 counterparts. In this study, we were able to deconvolve most atypical-grade meningiomas into a more indolent “grade 1-like” group and a more aggressive “grade 3-like” group, with concordant recurrence rates. These findings help further clarify the heterogeneity of this challenging disease, and support a shift towards molecular classification of meningiomas in clinical practice.

The World Health Organization^16^ currently recognizes molecular subclasses for some of the most aggressive brain tumors^17,18^, which have refined our predictions of treatment responses and clinical outcomes. Meningioma, the most prevalent type of adult brain tumor, are nevertheless categorized solely based on their histopathological appearances, a diagnostic heritage dating back almost eighty years^19^. Extremes of histological grades (mostly benign grade 1 and malignant grade 3) have relatively homogenous clinical behavior compared to grade 2 (or atypical) variants. There is a lot of uncertainty around the biology, clinical course and treatment response^20^ of atypical meningiomas. This is reflected by regular revisions in WHO definitions^21^, overlapping molecular signatures with adjacent grades^7,8^ and open questions about the benefits of chemotherapy^20,22^ and adjuvant radiation^23–25^. Defining subgroups of atypical meningiomas with homogenous biological and clinical properties may be critical to successfully resolving these questions, thereby improving prognostication and treatment for patients.

Given the relatively consistent biological features of benign grade 1 and malignant grade 3 meningiomas, we used them as genetic “hallmarks” around which to “deconvolve” the heterogeneous atypical class into “grade 1-like” and “grade 3-like” subgroups. Rather than relying on single gene markers, we used co-expression networks, a well-established system-based approach to analyze the gene expression arrays of 212 meningiomas from 6 independent cohorts (Supplementary table S1). The novelty of this systems-based approach is its sensitivity to small and cumulative signals from interacting genes within a biological cascade. This technique has been successful at defining gene expression patterns behind complex phenotypes in Huntington’s disease^14^, peripheral nerve regeneration^15^ or weight gain^11^. Though to our knowledge it has not been used to decipher molecular characteristics of atypical meningiomas.

## Results

We firstly established the gene expression profile that differentiates grade 1 from grade 3 meningiomas. Differential gene expression showed 1 up-regulated and 11 down-regulated genes (log_2_ fold change ≥ 1.5, p ≤ 0.0001) summarized in Figure 1A and Supplementary Table 2.

**Figure 1:**
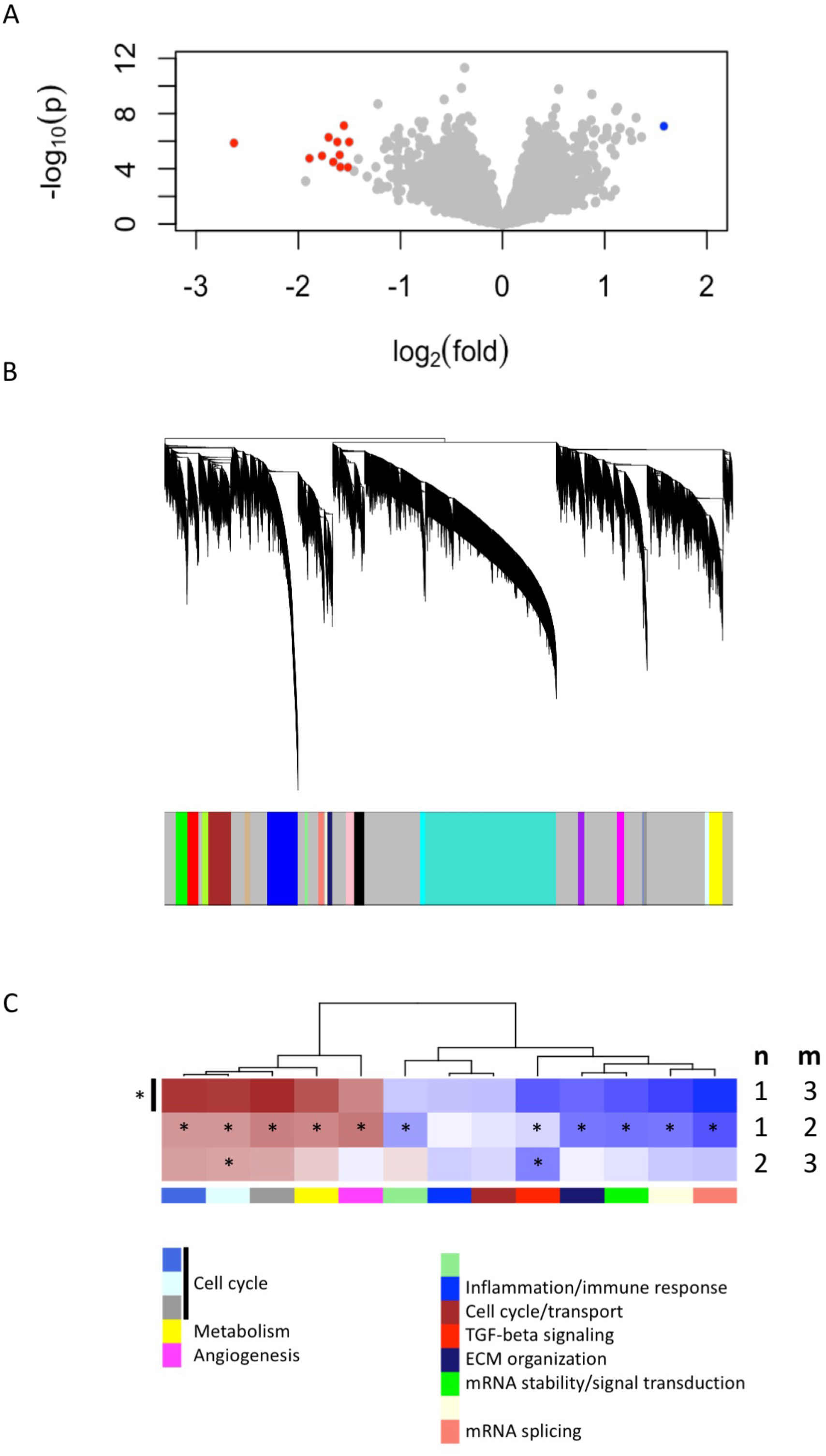
Gene expression signatures associated with meningioma grade. A: Differential gene expression between grades 3 and 1 meningiomas reveal 1 upregulated and 11 downregulated genes in grade 3 tumours (|log_2_(fold change) | ≥ 1.5, p<0.0001). B: Dendrogram of genes based on the topological overlap map, with the 20 gene modules represented as colors in the bar below. Grey represents unclassified genes. C: Plot of median module expression differences between grade (m) and grade (n). Only modules with significantly different expression between grades 1 and 3 are included (Mann Whitney p<0.05). Red indicates modules which are upregulated in grade (m), and darker shades indicate larger effect sizes. Notably, 11/13 modules are significantly different between grades 1 and 2 while 2/13 are also significantly different between grades 2 and 3. * p < 0.05 (Mann-Whitney).

We created another signature to distinguish grade 1 from grade 3 meningiomas using gene co-expression networks. This yielded 20 co-expressed gene modules (Figure 1B), of which 13 had expression levels that differed significantly between grades 1 and 3 (Mann-Whitney p<0.05, Figure 1C). A subset of these 13 were also significant between grades 1 and 2 and/or between grades 2 and 3 tumors, suggesting the intermediate biology of atypical meningiomas.

To find a genetic signature that best differentiates grades 1 and 3 tumors, we used two-centroid soft clustering and evaluated the resultant distribution of patients with a balanced sigmoidal cost function. An iterative feature selection approach was conducted using single genes and gene modules which were differentially expressed between grades 1 from 3. Notably, modules consistently yielded better performance (lower cost) than single genes (Figure 2A). The lowest cost was achieved with two modules as inputs; one of which contained 120 genes and is most associated with mRNA splicing while the other consisted of 98 genes and was involved in cell cycle. Using 80% membership probability as a cutoff, we reclassified 33 of 46 atypical meningiomas (72%) into a “grade 1-like” (14/46) and “grade 3-like” (19/46) subgroup of atypical meningiomas. (Figure 2B, Supplementary Figure 2). Recurrence rates were available for only a subset of cases (26/46) and were significantly higher in “grade 3-like” (8/11) compared to “grade 1-like” (0/9) subgroups (p <0.005).

**Figure 2:**
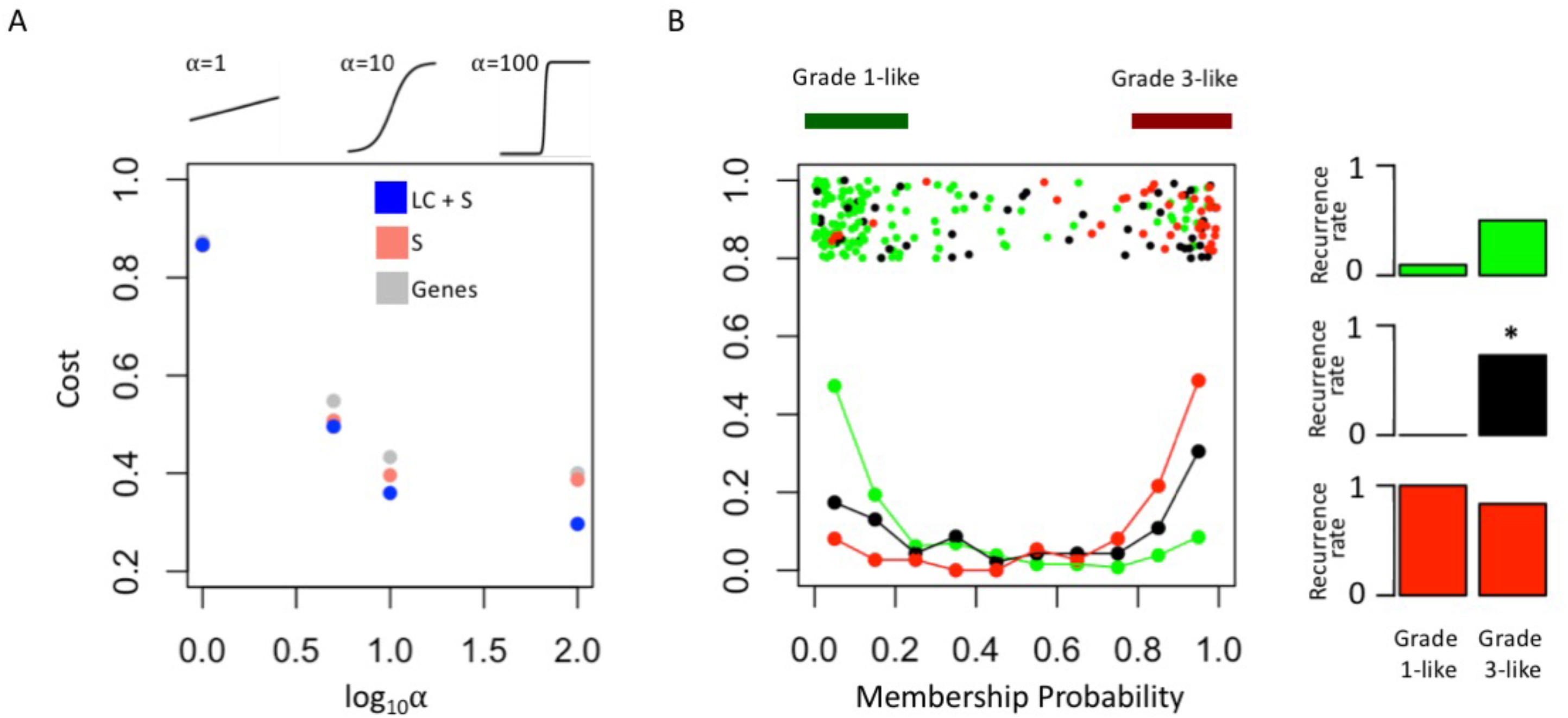
Optimized soft clustering reveals two subgroups of atypical meningiomas. A: Cost of multiple input configurations: “salmon”(S) and “lightcyan” (LC) modules in blue, “salmon” module alone and optimized differentially expressed genes in grey. Top inset depicts shape of sigmoid function with varied alphas. B: Summary graph of fuzzy C-means clustering best performing inputs (S + LC). The x-axis represents the probability of being in the grade-3 enriched cluster and y-axis represents the proportion of patients in each bin of 10%. Line graph component represents normalized frequency distribution of each histological grade (green = grade 1, black = grade 2, red = grade 3). Top jitter plot represents individual patients. Dark green and red bars above represent the 20 and 80% thresholding into grade 1-like and grade 3-like subgroups of atypical meningiomas. Recurrence rates are plotted on the right by grade (green, black, red) and subgroup (“grade 1-like” and “grade 3-like”). *Chi-square p<0.05.

Concordantly, there was no difference in recurrence rates between grade 1 and “grade 1-like” groups nor between grade 3 and “grade 3-like” groups.

Next, we verified the molecular identity of the newly detected subgroups of atypical meningiomas. Using a systematic comparison based on median module expression levels (Figure 3), we found concordance between the biology of our newly identified atypical subtypes with either grade 1 or grade 3 meningiomas (Figure 3A). Differential analysis also suggested that the overall biological separation between the newly described subgroups is similar to the separation between grades 1 and 3. This further lends to the validity of dividing atypical meningiomas into biologically homogenous subgroups which parallel existing grades.

**Figure 3:**
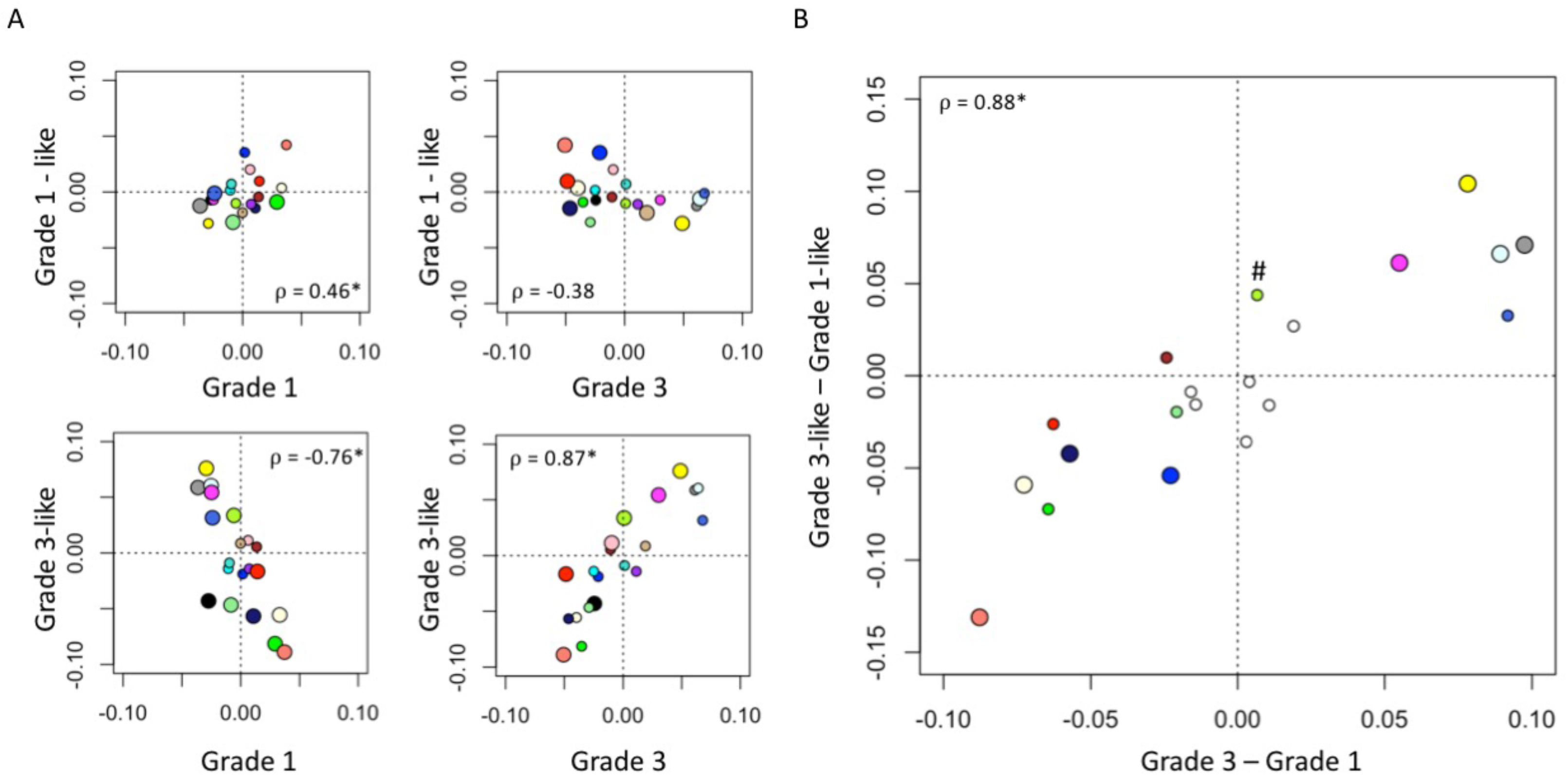
Molecular identity of newly described atypical meningioma subgroups. A: Scatter plot of median module expression. Larger circles indicate Mann Whitney p<0.05. Colors correspond to previously identified modules in Figure 1B. □=Pearson coefficient, *p<0.05. Note the positive correlation between the modules of grade 1 and “grade 1-like” and grade 3 and “grade 3-like” subtypes. B: Scatter plots of genetic separation between atypical subtypes as histological grades. The x-axis represents the difference in median module expression between grades 3 and 1, while the y axis represents the difference in median module expression between “grade 3-like” and “grade 1-like”. Large circles represent modules which are significantly different in both comparisons and empty circles indicate modules which are not significantly different in either. Of the remainder, 5/6 are significantly different between grades 3 and 1 only and 1/6 is significantly different between “grade 3-like” and “grade 1-like” (#).

## Discussion

The highly heterogeneous clinical behavior of atypical meningiomas suggests that histological criteria do not adequately capture it’s biology, thus motivating the discovery of molecular markers for better disease resolution. So far, molecular profiling of meningiomas has largely taken a monogenetic approach to marker discovery for aggressive phenotypes^9^. This has been fruitful in identifying recurrence mutations^7^ and transcripts^9^ linked to oncogenic cascades in meningiomas. These approaches however rely on differential gene expression to identify relevant molecular mechanisms and thereby remains limited in its ability to resolve small additive signal often relevant in tumor biology. Bypassing this problem, we applied gene co-expression networks to establish molecular signatures which identified indolent and aggressive subtypes of atypical meningiomas. Furthermore, majority of studies link genetic analysis to histological grade^7^, which does not capture disease biology for atypical meningiomas providing a 50-50 chance of recurrence. Epigenetic studies using conventional clustering have analyzed heterogeneity of meningiomas through all grades^8^ proposing new benign, intermediate and malignant methylation subclasses. “Intermediate” meningiomas are quoted a 20% chance of disease-free survival, more accurate than histology (50%). Our study focuses on the most heterogenous group, atypical meningiomas. We were able to separate subgroups with greater homogeneity compared to these preceding studies with 0% and 72,8% recurrence rates for grade 1-like and grade 3-like subgroups respectively. These findings also demonstrate the conceptual advantages of system-based approaches like co-expression networks over conventional techniques like differential gene expression and/or clustering. Our study has limitations: we analyze a single cohort, only a subset of samples have recurrence and follow-up times documented, on the other hand we provide a meta-analysis of 6 independent case-series which minimizes bias and counts towards the robustness of the findings.

Our findings help resolve the heterogeneity of atypical meningiomas by deconvolving into subgroups which are more homogenous then proposed prior studies. These homogenous subgroups may help predict clinical course, thus allowing for customized follow-up planning to manage resource intense investigations such serial imaging while optimizing patient care. Our results could also help guide recruitment protocols for future and ongoing clinical trials, which are currently limited by the uncertainty of clinical outcomes in atypical meningiomas^24^. The approach in this study lend to the utility of complex molecular signatures in augmenting histological diagnosis and resolving other heterogeneous and challenging diseases.

## Materials and methods

### Data collection and pre-processing

This study used open-source data from the repository *Gene Expression Omnibus (GEO)*^26^. Six studies with primary meningioma transcriptomics were included in the analysis, all of which had WHO grade annotated (Supplementary Table 1). These publicly available gene expression datasets were accessed Gene Expression Omnibus (GEO) under id’s: GSE10534, GSE77259, GSE54934, GSE43290 GSE16581 and GSE74385. For each study, the microarray data was backgrounded corrected, quantile normalized, and log-2 transformed using the *Affy*^27^ and *Limma*^28^ R packages for Affymetrix and Illumina platforms, respectively. After removing genes that were not common across these studies, the 6 studies were merged, scaled to a global mean and standard deviation of 0 and 1, respectively^29^, and batch-corrected using *ComBat*, a well-established empirical Bayes approach^30^. The resultant data matrix was used during all subsequent analysis.

### Differential gene expression analysis

Differential gene expression analysis was used to compare grades 1 and 3 meningiomas. In log2-transformed space, the fold change (FC) was computed by subtracting the mean expressions of each gene in grade 1 tumors from the corresponding mean expressions in grade 3 tumors. Genes with absolute log2-transformed FC ≥ 1.5 and p ≤ 0.0001 were considered significant.

### Co-expression networks and module detection

We used the well-established “Weighted Gene Correlation Network Analysis” (WGCNA) to detect “modules” of strongly co-expressed genes^13^. Per these previously described techniques, we first computed an “adjacency matrix” using soft-thresholded Pearson correlation between each gene pair. This was converted into a biologically-inspired topological overlap map (TOM), wherein pairwise gene similarities are derived from comparing their connectivity profiles^31^. Hierarchical clustering converted the TOM into a dendrogram, and a subsequent “dynamic” tree-cut^32^ served to identify gene modules. These modules were annotated with the *Database for Annotation, Visualization and Integrated Discovery* (DAVID)^33^, an open-source bioinformatics resource. Additionally, representative module “meta-genes” for each sample were computed as the first principal component of their constituent genes’ expression values. The utility of this approach was verified in our dataset by demonstrating that higher principle components capture a very small proportion of the overall variance (Supplementary Figure 1A) and showing that neither study batch nor sex cluster along the first principle component (Supplementary Figure 1B and 1C). This eliminates the possibility of batch effect or sex being drivers of our “meta-gene” values and confounding results. Differences in the expression levels of these “metagene” between grades was tested with a Mann-Whitney test, with a p≤0.05 considered significant.

### Feature selection and clustering

In order to better understand the heterogeneity of grade 2 meningiomas, we began by identifying genetic features able to best distinguish grades 1 and 3. Fuzzy C-means (FCM) clustering was applied to the set of all patients in our study and the resultant separation of grades 1 and 3 was established with a sigmoidal cost function that is balanced for differences in the prevalence of both grades:

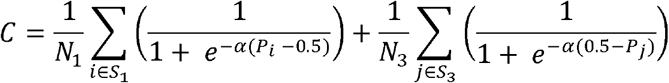

In the above equation, *N_1_* and *N_3_* are the number of grade 1 and grade 3 tumors, respectively; *S_1_* and *S_3_* are the sets of grade 1 and grade 3 tumors, respectively; *P_k_* is the FCM clustering-derived probability of patient *k* being in the grade 3-enriched cluster; and □ is a tunable hyperparameter. We used a two centroid model wherein cluster polarity was established by comparing the ratio of grade 3 to grade 1 tumours at both ends of the probability distribution (hard-thresholding at 80% probabilities).

Single genes and module “meta-genes” which were significantly different between grade 1 and grade 3 tumors served as input variables. Backwards elimination and forward selection were used for feature selection with model performance measured using the above cost function. Hyperparameter (□) values of 1, 5, 10, and 100 tested for all models. Once the separation of grades 1 and 3 was optimized, the probability distribution of grade 2 meningiomas within the same output was investigated. Atypical meningiomas with a probability ≥80% of being in the grade 1-enriched cluster were defined as “grade 1-like”, and those with a probability ≥80% of being in the grade 3-enriched cluster were defined as “grade 3-like”.

### Analysis of grade 2 subtypes

We first compared the recurrence rates of “grade 1-like” and “grade 3-like” meningiomas, and compared each to the rates of grade 1 and grade 3 tumors (Figure 2B, Supplementary Figure 2). Notably, only 114 of the 212 patients in our cohort have annotated recurrence. To investigate the degree of biological overlap between “grade 1-like” and grade 1 meningiomas, and similarly between “grade 3-like” and grade 3 meningiomas, we used the correlation between their module “meta-gene” expression levels (Figure 3A). In addition, we compared the biological separation between the newly described subtypes of atypical meningiomas to the separation of grades 1 and 3 by correlating their differential module expression levels (Figure 3B).

### Data analysis platforms

All computational work relied on the open-source computational platform R^34^, including packages *WGCNA*^13^, *ppclust*^35^, *Affy*^27^, *Limma*^28^, and *SVA*^36^.

## Supporting information

## Acknowledgements

The authors would like to thank Dr Anna Goldenberg and Miss Lauren Erdman for their suggestions regarding data analysis in this paper.

