## Supplementary material for "“Homogenous subgroups of atypical meningiomas defined using oncogenic signatures: basis for a new grading system?”"

**Supplementary Tables**

| GEO entry | Platform | Patients | Mean age  (SD) | N male  (%) | WHO grade (n)  I II III | Recurrence rate (%) |
| --- | --- | --- | --- | --- | --- | --- |
| GSE100534 | GPL6244 (Affymetrix) | 8 | N/A | 3 (37.5) | 6 1 1 | N/A |
| GSE77259 | GPL6244 (Affymetrix) | 14 | 54.1 (10.1) | 4 (28.6) | 10 4 0 | N/A |
| GSE54934 | GPL6244 (Affymetrix) | 22 | N/A | N/A | 20 2 0 | N/A |
| GSE43290 | GPL96 (Affymetrix) | 47 | 61.7 (15.0) | 13 (27.7) | 33 12 2 | 8/47 (17.0) |
| GSE16581 | GPL570 (Affymetrix) | 68 | 63.2 (14.7) | 25 (36.8) | 43 19 6 | 12/22 (54.5) |
| GSE74385 | GPL10558 (Illumina) | 53 | N/A | N/A | 17 8 28 | 22/45 (48.9) |
| Overall* |  | 212 | 61.7 (14.6) | 45 (32.8) | 129 46 37 | 42/114 (36.8) |

**Supplementary Table 1: Study demographics.**

*Of known values

**Supplementary Table 2: Differential gene expression in grade 3 versus grade 1 meningiomas**

| Gene Symbol | Gene name | log_2_(fold) | -log^10^(p) |
| --- | --- | --- | --- |
| PITX1 | Paired-like homeodomain 1 | +1.58 | 7.08 |
| EGFL6 | Epidermal growth factor like domain multiple 6 | -2.63 | 5.87 |
| LEPR | Leptin receptor | -1.89 | 4.76 |
| TMEM30B | Transmembrane protein 30B | -1.77 | 4.94 |
| TCEAL2 | Transcription elongation factor A like 2 | -1.70 | 6.28 |
| KCNMA1 | Potassium calcium-activated channel subfamily M alpha 1 | -1.66 | 4.49 |
| FGL2 | Fibrinogen-like protein 2 | -1.62 | 5.95 |
| ADAMTSL3 | A disintegrin and metalloproteinase with thrombospondin motifs like 3 | -1.60 | 5.00 |
| NDNF | Neuron-derived neurotrophic factor | -1.59 | 4.12 |
| ZC2HC1C | Zinc finger C2HC-type containing 1C | -1.55 | 7.12 |
| SLIT2 | Slit guidance ligand 2 | -1.51 | 4.10 |
| SERPINF1 | Serpin family F member 1 | -1.50 | 5.94 |
