## Supplementary figures and images for "“Homogenous subgroups of atypical meningiomas defined using oncogenic signatures: basis for a new grading system?”"

### Supplementary file 2

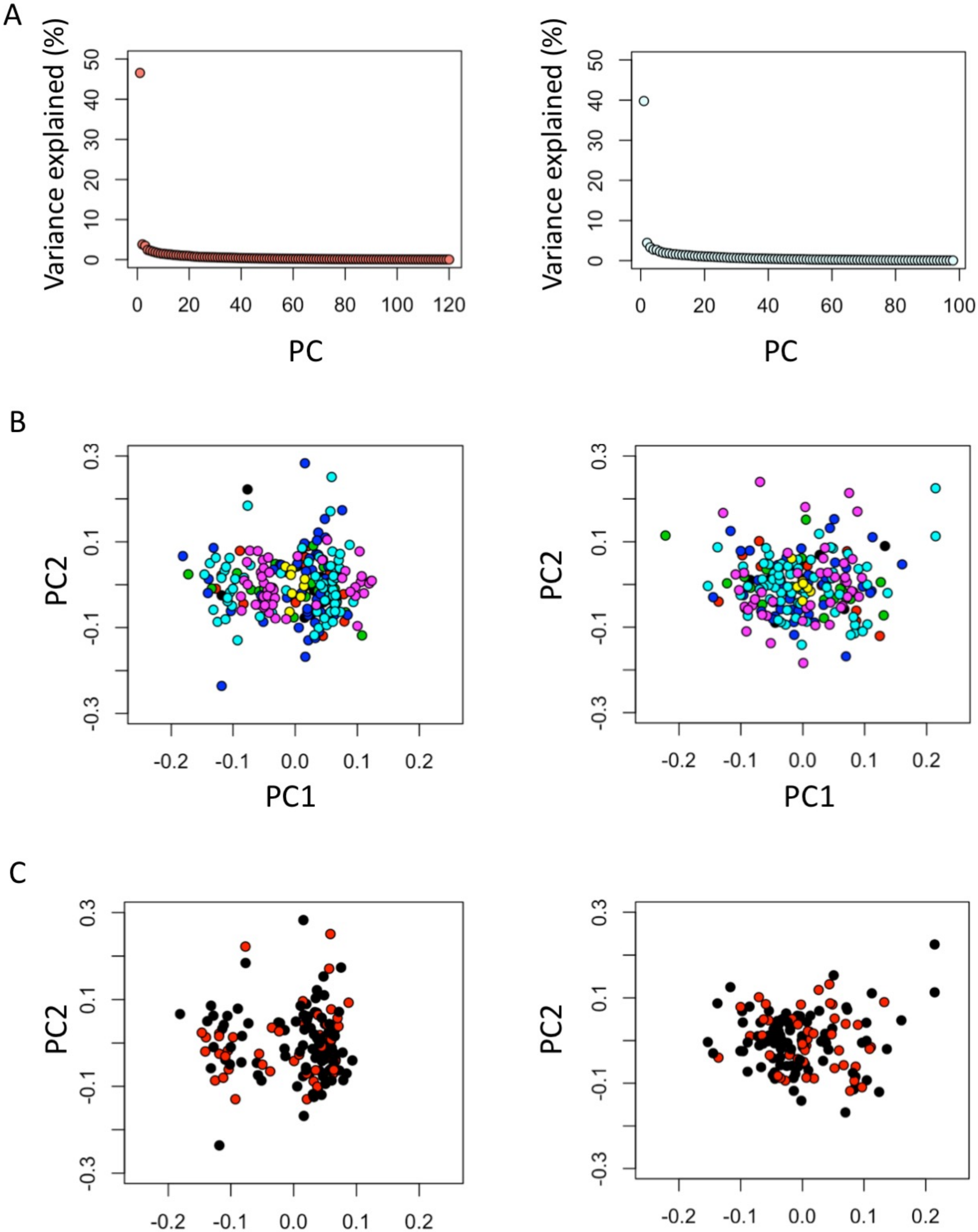


Supplement Figure 1

### Supplementary file 3

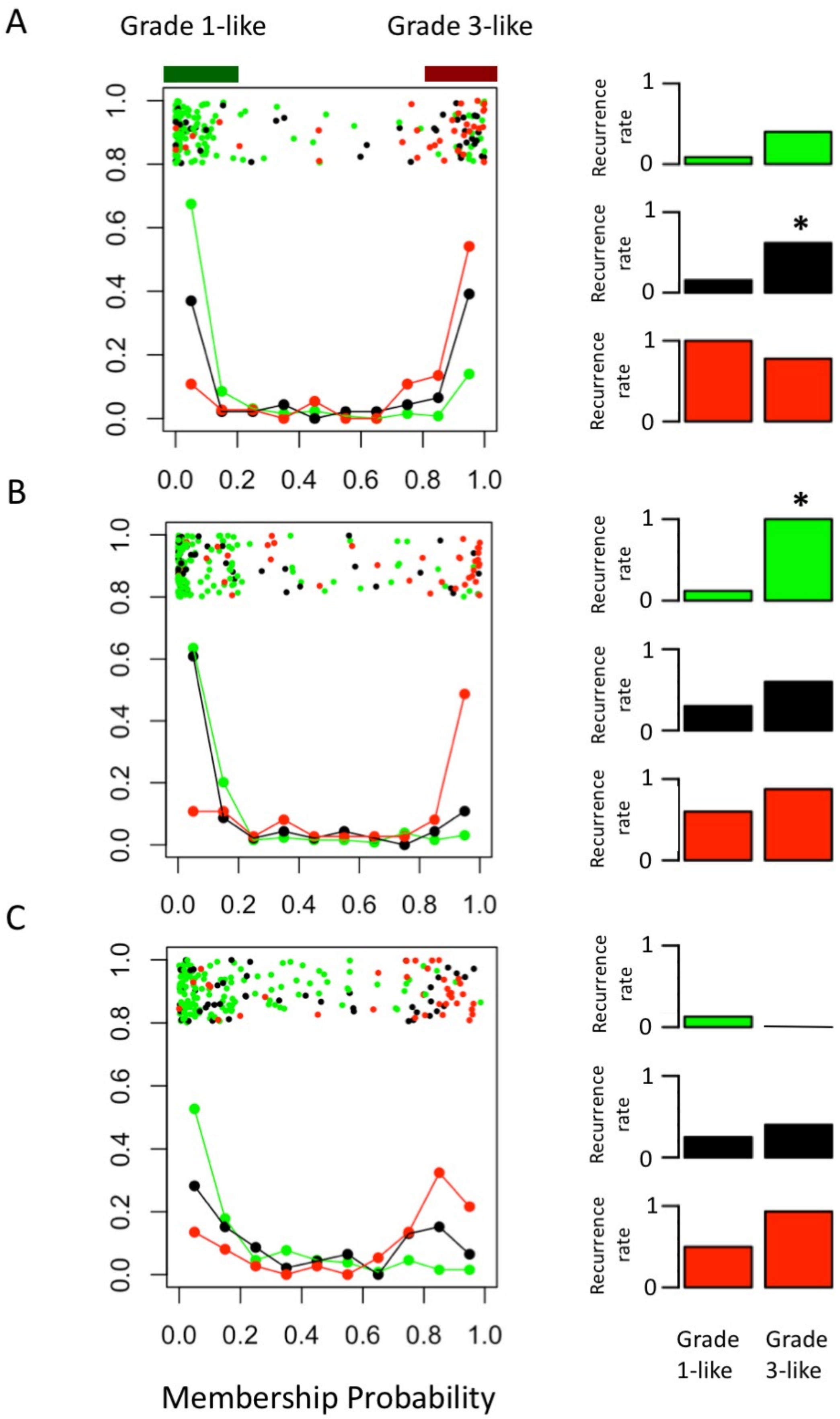


Supplement Figure 2
